# Assessing the Impact of Selective Attention on the Cortical Tracking of the Speech Envelope in the Delta and Theta Frequency Bands and How Musical Training Does (Not) Affect It

**DOI:** 10.1101/2024.08.01.606154

**Authors:** Alina Schüller, Annika Mücke, Jasmin Riegel, Tobias Reichenbach

## Abstract

Oral communication regularly takes place amidst background noise, requiring the ability to selectively attend to a target speech stream. Musical training has been shown to be beneficial for this task. Regarding the underlying neural mechanisms, recent studies showed that the speech envelope is tracked by neural activity in the auditory cortex, which plays a role in the neural processing of speech, including speech in noise. The neural tracking occurs predominantly in two frequency bands, the delta and the theta band. However, much regarding the specifics of these neural responses, as well as their modulation through musical training, still remain unclear. Here, we investigated the delta- and theta-band cortical tracking of the speech envelope of attended and ignored speech using magnetoencephalography (MEG) recordings. We thereby assessed both musicians and non-musicians to explore potential differences between these groups. The cortical speech tracking was quantified through source-reconstructing the MEG data and subsequently relating the speech envelope in a certain frequency band to the MEG data using linear models. We thereby found the theta-band tracking to be dominated by early responses with comparable magnitudes for attended and ignored speech, whereas the delta band tracking exhibited both earlier and later responses that were modulated by selective attention. Almost no significant differences emerged in the neural responses between musicians and non-musicians. Our findings show that only the speech tracking in the delta but not in the theta band contributes to selective attention, but that this mechanism is essentially unaffected by musical training.

## Introduction

Auditory selective attention refers to the ability of the human brain to segregate spatiotemporally overlapping speech streams into distinct auditory objects and to selectively attend one of them [1]. However, this ability requires significant cognitive resources and can be impeded by several factors, such as hearing impairment [2, 3], speech-related learning impairment [2] or age-related decline in the ability to attend to a target speaker [4, 5].

In contrast, musical training may prevent or even counteract difficulties in speech-in-noise perception [2, 3, 5, 6, 7]. Possible explanations for this hypothesis include that musical training can enhance brain plasticity [6] and functional connectivity [6, 8], increase the auditory working memory [9, 7, 10], and thus improve auditory fitness [2], especially when the training started at a young age [2, 11].

However, despite the importance of speech-in-noise comprehension for human oral communication and social life, the underlying neural processes, as well as the neural mechanisms leading to declined or improved abilities, are not yet fully understood. Recent investigations have employed non-invasive neuroimaging through electroencephalography (EEG) or magnetoencephalography (MEG) while participants listen to naturalistic speech in noise. Through subsequent statistical modeling, these recordings allow quantifying ongoing neural responses to repetitive, rhythmic aspects of speech stimuli, often referred to as neural speech tracking [1, 12, 13, 14].

Perhaps the most robust neural tracking emerges in response to the speech envelope, a signal in the low-frequency range, between 1-15 Hz, that traces the amplitude fluctuations of a speech signal [13, 15]. The neural tracking of the speech envelope does not simply reflect the bottom-up processing of the acoustic input, but is also influenced by higher cognitive processes, in particular by selective attention, which enhances the tracking [16, 1, 13]. Moreover, modulating the neural tracking through transcranial alternating current stimulation can impact and even enhance speech-in-noise comprehension [17, 18, 19, 20, 21].

Different functions have been attributed to cortical tracking in the theta band (4 - 8 Hz) and delta band (1 - 4 Hz) [22]. Theta-band tracking probably reflects the parsing of lower-level speech components, such as syllables and phonemes [22, 23, 24, 25], and reflects the acoustic clarity [26]. Delta-band tracking is associated with neural responses to words, reflects higher-level linguistic processing and can inform on speech comprehension [15, 26, 25]. Theta-band tracking has indeed been found to be more sensitive to stationary background noise, whilst delta band tracking seems to be robust as long as the speech stimulus is still intelligible [15, 27]. However, the specific contributions of delta- and theta-band tracking to selective attention have not yet been fully clarified.

The neural specialization that may allow musicians to achieve better speech-in-noise comprehension has been in-vestigated through non-invasive neuroimaging as well, mostly focusing on the evaluation of short auditory stimuli [28, 9, 29, 30]. These investigations found, in particular, enhanced subcortical responses of people with musical training [31, 32, 29], as well as a larger right-hemispheric recruitment of neural resources in the auditory cortex [33].

However, there has only been very little work investigating the impact of musical training on cortical speech tracking. As a notable exception, Puschmann et al. (2019) found that in musically-trained individuals, attentional modulation of the cortical speech tracking was less pronounced, suggesting that the ignored stream is represented stronger in cortical activity [34]. However, they did not investigate delta and theta band cortical tracking separately.

Here, we sought to differentiate the modulation of the neural tracking in the delta and in the theta band through selective attention, as well as how these individual responses may be shaped by musicianship. We hypothesized that attentional modulation only acts on the higher-level processing associated to the delta band, but not on the lower-level processing in the theta band, since the attentional effects have been observed comparatively late, at delays of 140 ms and later [35]. We further hypothesized that the attentional modulation of the delta-band tracking is more pronounced in musicians, contributing to an enhanced behavioral performance in speech-in-noise listening.

## Materials and Methods

### Experimental Design

We used an MEG dataset from 52 participants (26 female, 26 male) aged 24.1 *±* 3.1 years that was acquired in the scope of two of our previous studies [36, 37]. All participants were right-handed and native German speakers. They had no history of neurological disease or hearing impairment. The study was approved by the ethics board of the University Hospital Erlangen (registration no. 22-361-S).

Participants listened to two German audiobooks simultaneously, both of which were narrated by male speakers (Figure 1). Two audiobooks were used alternatingly as *attended audiobooks*, which the participants were instructed to focus on. The first attended audiobook was *‘Frau Ella’* written by Florian Beckerhoff and read by Peter Jordan (*Speaker 1*). The second attended audiobook was *‘Den Hund überleben’* written by Stefan Hornbach and read by Pascal Houdus (*Speaker 2*). Simultaneously, an incoherent excerpt from an unrelated audiobook read by the other speaker, respectively, was presented as *ignored audiobook*. The first ignored audiobook was thus read by *Speaker 2*, Pascal Houdus, and was titled *‘Looking for Hope’* by Colleen Hoover (translated into German by Katarina Ganslandt). The second ignored audiobook was *‘Darum’* by Daniel Glattauer and read by Speaker 1, Peter Jordan. All of the employed audiobooks were published by *Hörbuch Hamburg* and are available in stores. Speaker 1 read with a mean word frequency in the first attended audio of 3.6 Hz with an average syllable frequency of 5.8 Hz. In the second attended audio, Speaker 2 showed comparable speech characteristics with a mean word frequency of 3.7 Hz and a mean syllable frequency of 5.7 Hz. The audio data, as well as the MEG recordings, will be published upon submission.

**Figure 1:**
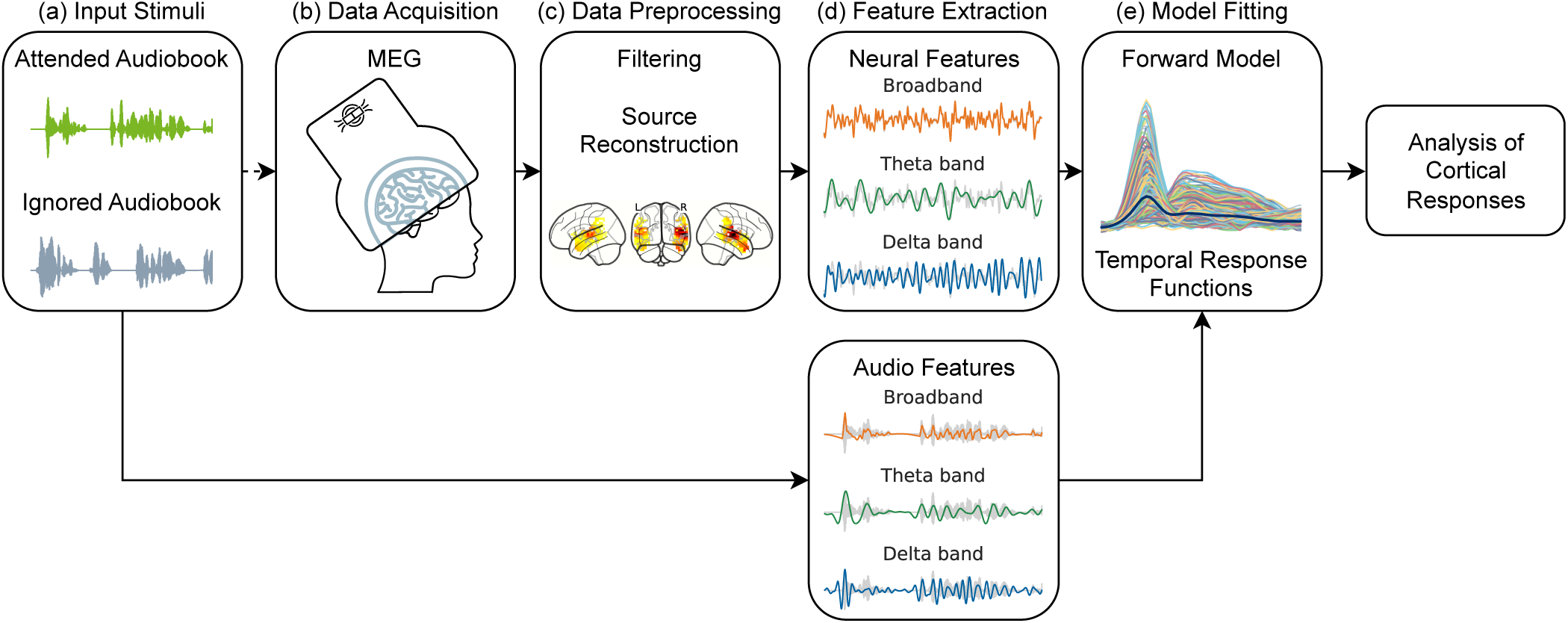
Overview of the experimental setup and the data processing pipeline. (a) Participants were presented with two audiobooks while attending one and ignoring the other. (b) Simultaneously, MEG was measured. (c) Based on the acquired data, source reconstruction with focus on the auditory cortex was performed. (d) From the source-level data, the frequency bands of interest were extracted using bandpass filtering. The corresponding audio features were extracted from the acoustic envelopes of the ignored and attended audio, respectively. (e) Lastly, temporal response functions (TRFs) were calculated by training a forward model to estimate the neural features from the audio input features. The resulting TRF magnitudes were compared between the attended and ignored condition, as well as between musicians and non-musicians.

The experiment was split into ten trials. Each trial consisted of listening to one chapter from one of the attended audios. Simultaneously, a random duration-matched excerpt from the corresponding ignored audio stream was presented. Once the audiobook chapter ended, participants had to answer three single-choice listening comprehension questions about this chapter to ensure that the participants did indeed focus on the attended audio. After each trial, the attended audio, and thus the attended and ignored speakers, were switched. This was repeated until the first five chapters of both attended audios were presented. The chapters had varying lengths between 3–5 minutes, resulting in a total recording duration of 37 minutes. The audios were presented diotically at equal sound pressure levels of 67 db SPL(A).

MEG data were acquired with the participants in a supine position and with their eyes focused on a fixation cross displayed on a screen above. Their head was placed in a 248-channel MEG system (*4D Neuroimaging*, San Diego, CA, USA). Before the start of the measurement, the head shape of each participant was captured using a digitizer (*Polhemus*, Colchester, Vermont, Canada). Additionally, the head position relative to the MEG system was recorded using five integrated head-position indicator coils. MEG data was recorded with a sampling frequency of 1017.25 Hz. During online processing, an analog bandpass filter (1 - 200 Hz) and a noise reduction algorithm (*4D Neuroimaging*, San Diego, CA, USA) using 23 reference sensors were applied to eliminate environmental noise.

The audio stimuli were presented to the participants using a customized setup that was validated and described in more detail in previous work [38]. In brief, it consisted of two flexible tubes that were connected to loudspeakers and led into the magnetically-shielded MEG chamber, connecting to earphones that the participants wore. Simultaneously, the presented audio signal was fed as an additional input channel to the MEG data logger. This enabled the synchronization of MEG recordings and the auditory stimuli with a 1 ms precision.

### Assessment of musical training

Participants were assigned to three groups based on their history of musical training, musicians (M), non-musicians (NM), and neutral participants (N). This categorization was performed using previously established criteria [37], namely starting age of playing an instrument, total years of undergoing musical training, and whether they were currently practising.

To be classified as a musician, a participant had to start playing before the age of seven for a total period of ten or more years. Furthermore, they had to be regularly play an instrument at the time of the study. On the other hand, participants were classified as a non-musician if they did not undergo musical training any longer than three years in sum and only started aged older than seven. Participants who did not fit in either of these categories were assigned to a third, neutral group.

Based on these criteria, we acquired data from 25 non-musicians (12 female, 13 male, aged 24.5 *±* 3.4 years), 18 musicians (10 female, 8 male, aged 24.1 *±* 3.1 years), and 9 neutral participants (4 female, 5 male, aged 24.2 *±* 3.9 years). Analysis that disregarded musical training was conducted using the data from all 52 participants. For the comparison between musicians and non-musicians, participants from the neutral group were excluded, resulting in data from 43 participants. This number of participants is in line with previous studies that investigated speech processing with regards to the musicality of a subject [29, 31, 32].

### Data Analysis

Data processing and analysis were performed in Python using the libraries *MNE* [39], *SciPy* [40], and *statsmodels* [41].

### MEG Data Preprocessing

First, the acquired MEG recordings were cut to the intervals of interest where an acoustic stimulus was present. These pieces were then concatenated.

The MEG data were further processed with a notch filter (50 Hz) to remove powerline interference. Afterwards, the frequency range of interest was extracted using a Butterworth bandpass filter (1 - 20 Hz, order *n* = 4). This filter was applied forward and backward to prevent introducing a filter shift. To increase computational efficiency, the filtered data was downsampled to 100 Hz.

For source reconstruction, we used the average MRI brain template ‘fsaverage’ provided by the *FreeSurfer* software package [42]. The computational steps were performed using the Python package *MNE* [39]. The ‘fsaverage’ template was aligned to the participant-specific head positioning and head shape that was collected before each measurement.

Next, a volume source space consisting of an equidistant grid of sources was created. As our analysis focused on auditory processes, sources were exclusively created in the auditory cortex and its surrounding region, including the middle temporal gyrus, the transverse temporal gyrus, the superior temporal gyrus and its banks, the supramarginal gyrus, and the insular lobe, in both the left and right hemisphere. For the subcortical segmentation and cortical parcellation of the brain regions within the volume, the ‘aparc+aseg’ template from *FreeSurfer* was employed. Sources were selected to have a distance of 5 mm between one another, resulting in 525 distinct source points.

Using the head model and the source space, a forward solution was calculated estimating the magnetic field strength at each MEG channel produced by the sources. To account for the different types of tissue in the brain and their varying conductivities, the boundary element method (BEM) model provided by *FreeSurfer* for the ‘fsaverage’ template was employed. The resulting leadfield matrix characterizes the sensitivity of each MEG channel to the activity of each of the selected sources.

A spatial filter was then applied to reconstruct the source activity from the sensor-level data and the estimated leadfields. Here, a linearly constrained minimum variance (LCMV) beamformer [43] was employed to estimate the activation of each source independently whilst deducting environmental noise measured in the empty MEG chamber.

For analyzing neural tracking in the delta and theta band, the data was further filtered in the corresponding frequency range. To this end, we employed a forward-backward Butterworth bandpass filter (order *n* = 4) in the range of 1 - 4 Hz and 4 - 8 Hz, respectively.

### Speech envelope

The audio files had a sampling frequency of 44.1 kHz. To retrieve the acoustic speech envelope, the analytic repre-sentation was obtained by applying a Hilbert transform. The magnitude of this analytic signal was then calculated, as it represents the instantaneous amplitude and hence the envelope of this signal.

To analyze the neural tracking of the audio, we filtered the speech envelope in three different frequency bands. On the one hand, a broad range of frequencies ranging from 1 - 20 Hz was extracted, which is referred to as *broadband* feature in the following. On the other hand, two narrower frequency bands were analyzed as well, namely the *delta band* (1 - 4 Hz) and the *theta band* (4 - 8 Hz). The envelope was then filtered within the desired frequency bands using a Butterworth bandpass filter (order *n* = 4) applied forward and backward to prevent filtering delays using the *SciPy* function sosfiltfilt [40]. The resulting features were then resampled to 100 Hz to match the sampling rate of the preprocessed, source-level MEG data.

### Temporal Response Functions (TRFs)

To analyze the neural tracking of the extracted audio features, we determined the relation between them and the neural data. Using the extracted audio features as input, we calculated forward models to predict the corresponding neural responses to both the attended and ignored audio and for each frequency band.

For this purpose, a linear model was trained that predicted the neural response 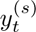 for every point *t* in time at every neural source *s* based on a linear combination of acoustic feature values *x_t−τ_* covering the time interval [*t − τ*_max_*, t − τ*_min_]. For each time delay *τ*, a coefficient 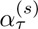 was estimated. *τ_min_* was chosen to be negative to verify the plausibility of the resulting model weights and to gain information about the noise level present in the TRF. As not all variation in the neural signal can be explained by the auditory input, a residual 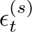 remains for every investigated time lag:

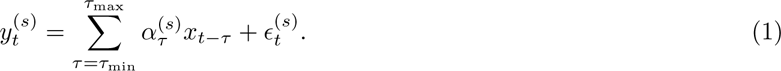

The coefficients 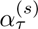 are referred to as the TRF of the source point *s*. The relative magnitudes of these coefficients characterize to which extent each delay contributes to the neural signal, and thus, how strong the neural response to the auditory input is after a certain time lag. Local maxima in the magnitudes of the TRFs indicate that neural responses are particularly pronounced at these time delays.

However, unconstrained linear regression is not the optimal approach for speech features as an input signal, as it likely contains temporal correlations causing numerical instability when fitting the model [44]. Furthermore, due to the high amounts of input data and parameters, the model is prone to overfitting. To address these issues and thus retrieve more robust and reliable results, we employed ridge regression [45, 46, 47, 36]. The TRF coefficients ***α****^(s)^* ∈ ℝ^(*τ*_max_−*τ*_min_)×1^ were accordingly approximated as

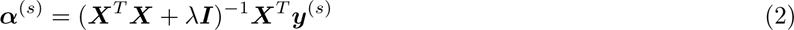

where ***X*** ∈ ℝ^Δ*t*×(*τ*_max_−*τ*_min_)^ contains the speech envelope information over the entire stimulus length Δ*t*, ***y****^(s)^* ∈ ℝ*^T ×^*^1^ describes the neural recording at source *s*, *λ* is a predefined regularization parameter, and ***I*** refers to the identity matrix. A Python implementation of this TRF coefficient estimation method that was previously developed and utilized by our group [46, 47] was used. To retrieve meaningful results, all speech features and neural features were scaled using z-score normalization.

To select the regularization parameter, a subject-wise five-fold cross-validation testing 12 parameters ranging from 10*^−^*^5^ to 10^5^ was applied. The respective optimal *λ* was calculated based on the prediction that yielded the highest correlation with the measured neural signal. Due to similar results across all participants, the mean optimal *λ* from the search space was used for all TRF coefficients per investigated frequency band to ensure comparability of the results. The resulting regularization parameters are listed in Table 1.

**Table 1:**
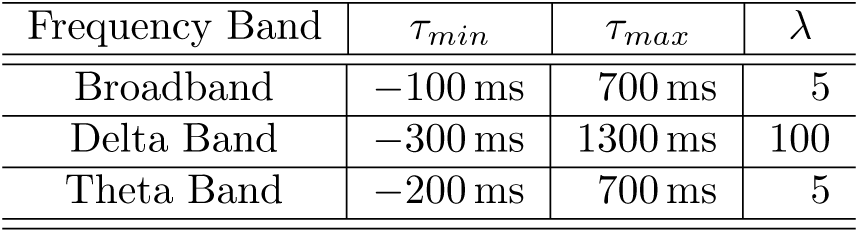
Overview of the parameters employed for computing the TRFs.

| Frequency Band | $\tau_{min}$ | $\tau_{max}$ | $\lambda$ |
| --- | --- | --- | --- |
| Broadband | −100 ms | 700 ms | 5 |
| Delta Band | −300 ms | 1300 ms | 100 |
| Theta Band | −200 ms | 700 ms | 5 |

The time lag intervals were selected based on visual inspection of the TRF estimates and are depicted for all frequency bands in Table 1. A step size of 10 ms between each time step was used, which corresponds to the sampling frequency of 100 Hz of the employed features.

### TRF Peak Extraction

Distinct TRF coefficients were estimated for all participants and the two listening modes *attended* and *ignored*. Our main focus was to identify the most prominent responses and corresponding time lags and compare their intensities across different groups and conditions. Thus, to simplify the subsequent analysis, only the magnitudes of the individual TRFs were further regarded. The TRF magnitudes were min-max normalized at the population level.

To determine the magnitude and average latency of the distinct peaks in the TRFs, the average TRF magnitudes were investigated at the population level. They were calculated by averaging the TRF magnitudes over all source points, all participants, and both presented speakers, for both attended and ignored listening modes. Based on these participants’ average responses, the most prominent peaks and their latencies were identified for each frequency band and listening mode.

As the TRF peak latencies differed from one subject to another, we found that information loss occurred when individual peak magnitude values were retrieved by simply considering participants’ average peak latencies. Thus, we also extracted a set of individual TRF peak latencies and magnitudes from the source-averaged TRF magnitudes for each individual participant. To avoid distorting the results with very early or late outlier peaks that may appear in individual TRFs due to noise or artifacts, we defined a fixed search interval around each average peak per frequency band. For every peak type, all local maxima were identified and the respective maximum closest to the average was selected as the individual peak. If no local maximum was found in the search interval, the individual TRF was excluded from the analysis of the respective peak.

### Attentional Modulation of Cortical Responses

To investigate the impact of attention on the cortical responses, TRF magnitudes were investigated on the single-subject level. For each participant, cortical responses to the attended and ignored audios were retrieved, using the envelope from the attended and ignored streams as input features, respectively. Thus, the two listening modes *attended* and *ignored* were compared to evaluate differences in cortical tracking of the respective speech envelopes. TRF magnitudes were obtained by averaging over all sources. From these subject-level TRFs, individual TRF peaks characterized by peak amplitudes and peak latencies were extracted as described above and compared between listening modes.

To quantify the attentional modulation of the cortical responses for each participant and compare this measure between the non-musician and musician groups, an attentional modulation index *AI* was calculated for each participant, speaker, frequency band, and TRF peak *p* as

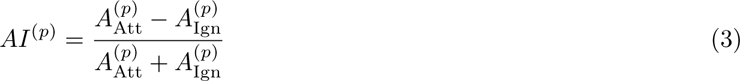

where 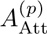 describes the individual peak amplitude at peak *p* for the attended and 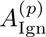 for the ignored audio feature, respectively. Values of *AI* close to zero indicate similar response strengths to the ignored and attended stream and hence, no attentional modulation. In contrast, positive values of *AI* suggest the existence of attentional modulation by attenuation of the ignored stream or amplification of the attended stream, whereas negative values of *AI* imply the opposite effect, i.e., a stronger cortical representation of the ignored than the attended stream.

### Statistical Analysis

All statistical tests were performed using the Python package *statsmodels* [41] and the *stats*-module from the Python package SciPy [40]. The significance level was set to *p <* 0.05 for all tests.

### Significance of Peaks in Population-Level TRFs

The population-level TRFs for both the attended and the ignored listening mode were obtained by averaging over the magnitudes of the TRF coefficient from all participants and sources. The intervals in which these population-level TRF magnitudes were significantly different from noise were determined using a bootstrapping approach that compared the magnitudes to a distribution of noise models. Participant- and source-specific noise models were generated by fitting the ridge regression model on a reversed version of the auditory input features. Therefore, the noise models did not contain any meaningful relationship between the input feature and the predicted neural output.

The resulting individual noise model weights were randomly shuffled across time lags, channels, and participants and then averaged in the same manner as the individual TRFs to generate the participants’ average noise over all regarded time lags. This procedure was repeated 10.000 times to obtain a noise model distribution. Using this distribution, empirical *p*-values were calculated for each TRF and time lag based on the amount of noise-level values with a lower magnitude than the investigated actual TRF magnitude. To account for multiple comparisons across all time lags, the resulting *p*-values were revised using the Bonferroni correction.

### Significance of Lateralization

To determine how the spatial distribution of cortical responses to natural speech differs between earlier and later time delays and across the investigated frequency bands, source-specific average TRFs were calculated. This was achieved by averaging the source-level TRFs over all participants and attended audios. Only the attended audios were incorporated in this analysis, as we wanted to analyze the spatial distribution of responses independent of possible top-down effects introduced by attentional modulation. As a result, the cortical responses to the attended speech envelope for each of the 525 source points in the area of the auditory cortex were retrieved. Snapshots from the source-level TRFs were extracted at each of the previously determined average response peak latencies. The momentary spatial distribution of intensities in the source space at these time lags was visualized through brain plots.

To assess whether potential differences in these responses occurring between sources in the right and left hemispheres were statistically significant, a two-sided Mann-Whitney-U test was performed. This test was undertaken with the TRF magnitudes at the latencies of the population-level TRF peaks in all frequency bands. To correct for multiple comparisons for the different peaks within a frequency band, the Bonferroni correction was applied.

To further quantify the extent of lateralization at the response peaks yielding significant differences between both hemispheres, a participants’ average lateralization index *LI*^(*p*)^ for every peak *p* and frequency band was calculated as already applied similarly in previous literature [48]:

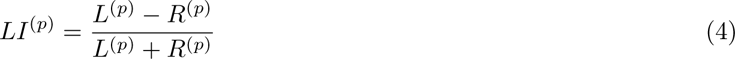

where *L*^(*p*)^ and *R*^(*p*)^ describe the sum of peak magnitudes from sources in the left and right hemispheres, respectively. Thus, a negative *LI*^(*p*)^ value indicates right-lateralized neural activity, whereas a positive *LI*^(*p*)^ value implies a left-lateralized neural processing.

### Significance of Attentional Modulation

The degree to which attended and ignored speech were tracked differently in the auditory cortex was assessed at the population level using different characteristics of the TRF peaks, such as peak amplitude, peak latency, and the attentional modulation index. When a certain TRF peak was not present in one listening mode for a particular participant, e.g., for the ignored audio, the respective peak was also excluded for the respective other mode, e.g., the attended audio, to allow a mode-wise comparison within participants. Statistical significance of differences in these metrics within the same group of participants, i.e., when comparing attended and ignored speech responses for all participants, were assessed using a two-sided pairwise t-test if the underlying data of the respective characteristic was normally distributed. Otherwise, a two-sided Wilcoxon signed-rank test was applied. The normality of the analyzed samples was determined using the Shapiro-Wilk test.

When comparing characteristics between musicians and non-musicians, a two-sided unpaired t-test was utilized for normally distributed characteristics. Alternatively, for samples that did not follow a normal distribution, a two-sided Mann-Whitney-U test was utilized. For all tests within a frequency band, the Bonferroni correction was applied to compensate for multiple comparisons across the different TRF peaks.

### Relationship of Listening Comprehension and Cortical Response Strength

During the study, participants answered single-choice listening comprehension questions regarding the attended audio stream. The percentage of correct answers per participant and attended audiobook was captured as a behavioral metric to quantify the performance of the individual participant. To relate the cortical response to this behavioral measure, Pearson’s correlation coefficients were calculated between the comprehension score and each TRF peak per feature and listening mode. The Bonferroni correction was used to adjust for multiple comparisons between different TRF peaks for all correlations within a frequency band. Furthermore, to evaluate the listening performance of musicians and non-musicians on a purely behavioral level, a two-sided unpaired t-test was performed to compare the comprehension scores of the two groups.

### Relationship of Musicality Features and Cortical Response Strength

As detailed above, participants were asked about different aspects of their musical training: their starting age of playing an instrument, total years of undergoing musical training, amount of current training, and the total number of instruments played. (Note that only the first three aspects were used to classify the participants into the three categories.)

Using a linear regression model, we assessed if the TRF peak characteristics related to these reported musicality aspects. A regression model was estimated for each peak using ordinary least squares. Due to the large number of estimates, this analysis was only performed for the delta and theta bands. All independent and dependent variables were scaled beforehand using z-score normalization. The musicality aspects served as independent variables, whereas the peak values were used as the dependent predicted features.

## Results

### Temporal and Spatial Distribution of Cortical Responses to Attended Speech

As a first step, we assessed the temporal and spatial distribution of the neural speech tracking in the three employed frequency bands at the population level (Figure 2). For the broadband responses, the average magnitude of the TRFs exhibited peaks at the time lags 110 ms and 240 ms (Figure 2a). In the delta band, peak neural responses emerged at the time lags 100 ms, 270 ms, and 540 ms. The cortical tracking in the theta band led to a main peak at 130 ms. The symmetrical sidelobes around this peak are caused by the narrow bandwidth of the speech feature and were therefore not regarded as independent peaks.

**Figure 2:**
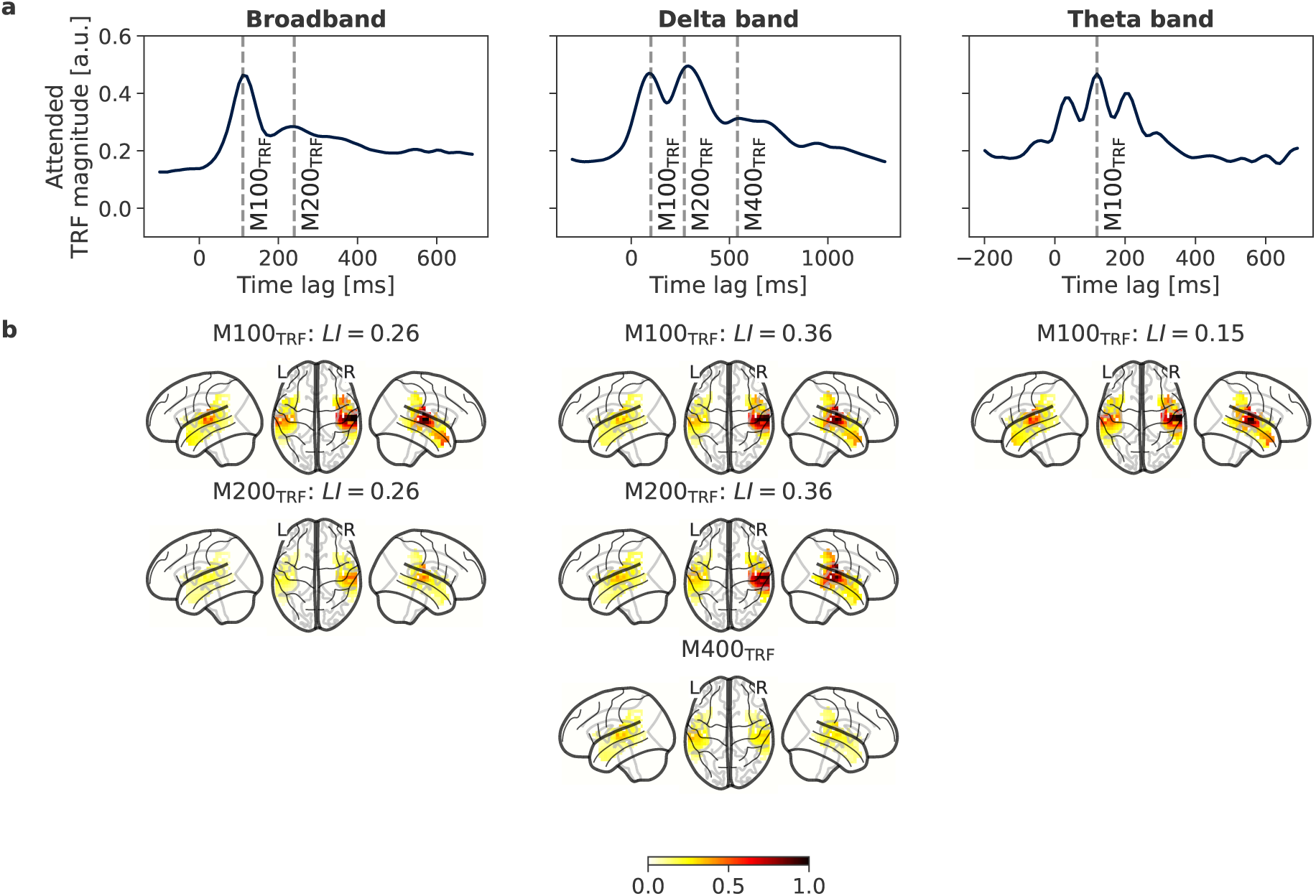
Population-level magnitudes of the averaged temporal response functions (TRFs) of attended speech and their spatial distri-bution for the distinct frequency bands. Panel (a) shows the mean magnitudes of the TRFs across all source channels and participants for attended speech for each investigated frequency band. Based on these functions, the time lags at which cortical responses were maximal (dotted vertical lines) were extracted and assigned to one of the response types M100_TRF_, M200_TRF_, or M400_TRF_. Panel (b) exhibits the distribution of TRF magnitudes at these maximum response times in the source volume ranging from low activation (yellow) to high activation (dark red). For brainplots showing significant differences in activations of right and left hemispheres, the respective lateralization indices (LI) are reported. The colored area corresponds to the investigated source region of interest

In the following, responses peaking around 100 ms and 200 ms are referred to as M100_TRF_ and M200_TRF_, respectively. Furthermore, the late TRF response peak observed in the delta band is referred to as M400_TRF_ in analogy to the N400 component in event-related potentials [49]. We thus found a M100_TRF_ in all three frequency bands, a M200_TRF_ in the broadband and the delta band, and a M400_TRF_ solely in the delta band.

M100_TRF_ showed a highly significant right-lateralization in all three frequency bands, with lateralization indices *LI* = 0.26 (*p* = 6e*−*19) in the broadband, *LI* = 0.36 (*p* = 1e*−*40) in the delta band, and *LI* = 0.15 (*p* = 2e*−*11) in the theta band (Figure 2b). A similar degree of right lateralization was observed for M200_TRF_ with *LI* = 0.26 (*p* = 1e*−*31) in the broadband, and *LI* = 0.36 (*p* = 5e*−*46) in the delta band. In contrast, M400_TRF_ showed a bilateral distribution of activations (*LI* = 0.01, *p* = 0.99).

### Attentional Modulation of Cortical Responses

In the next analysis step, TRF responses to the ignored audios were compared to the attended responses at the population level (Figure 3). For all three frequency bands, the TRF magnitudes were significantly higher than those of the noise model for the majority of the analyzed positive time lags.

**Figure 3:**
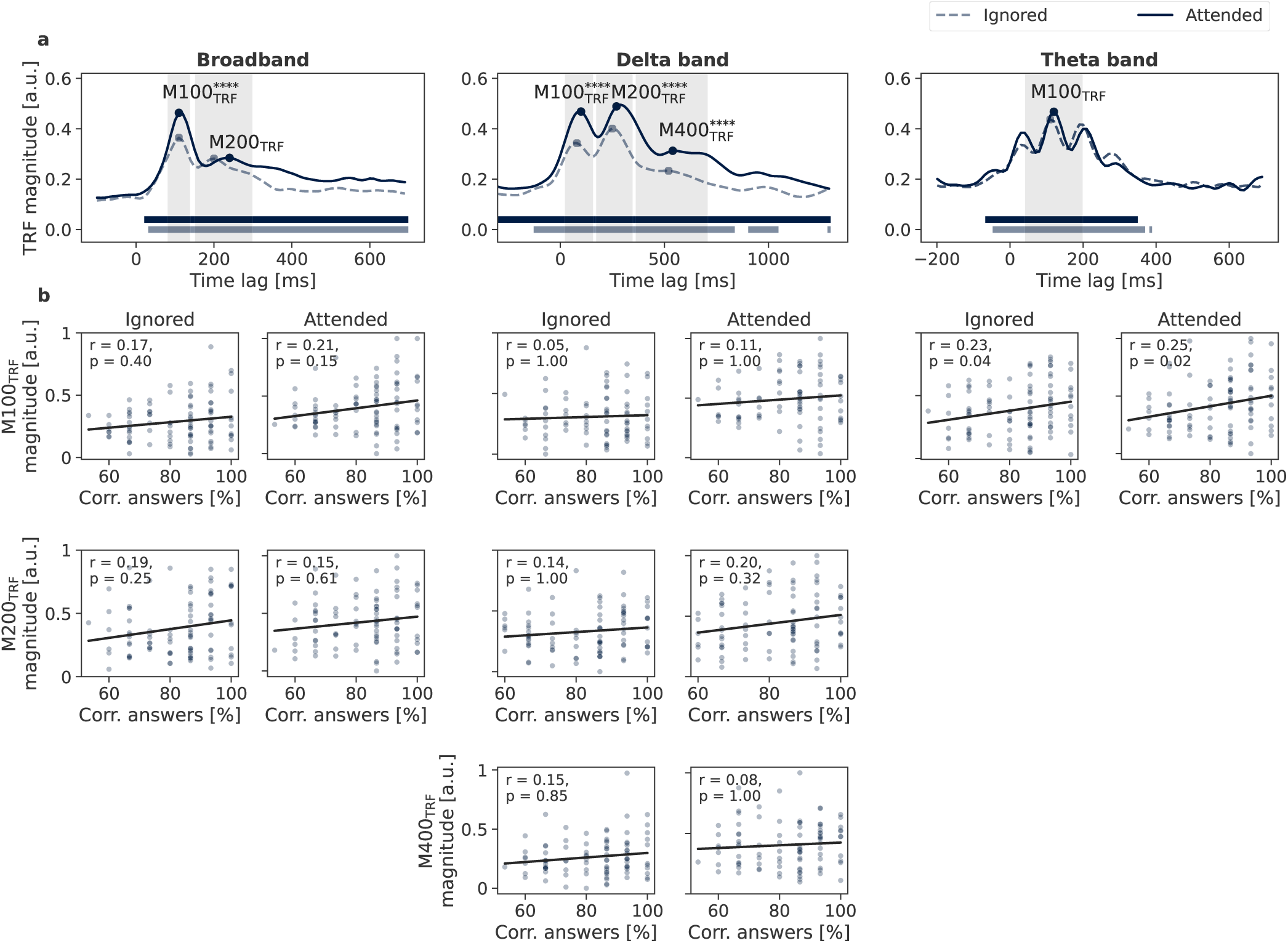
Population-level TRF magnitudes in the attended and ignored condition, as well as their relation to the comprehension scores. (a) The TRF magnitudes in the attended condition (black solid) are larger than those in the ignored condition (grey dashed) for the broadband and the delta-band response, but not in the theta band. The horizontal bars indicate the areas where the TRF magnitudes significantly differ from the noise models. Peaks with significant differences in amplitude between the attended and ignored condition are marked with asterisks indicating their significance level (*, 0.01 *≤ p <* 0.05; **, 1e*−*3 *≤ p <* 0.01, ***, 1e*−*4 *≤ p <* 1e*−*3, ****, *p <* 1e*−*4). The search areas for the participant-wise TRF response peaks are shaded in grey. (b) The individual magnitudes at each peak were extracted per subject and related to the percentage of correct comprehension question answers through Pearson’s correlation coefficient *r*. A significant positive correlation between response magnitude and comprehension was found in the theta band, both for the attended and the ignored condition

Analogous to the attended TRFs, the ignored TRFs in the broadband condition exhibited both M100_TRF_ (time lag 110 ms) and M200_TRF_ (time lag 200 ms). The same responses were also visible in the ignored TRFs in the delta band, with M100_TRF_ and M200_TRF_ occurring with average delays of 80 ms and 250 ms, respectively. Additionally, weak M400_TRF_ response was extracted at 520 ms to allow a direct comparison with the attended responses. As for the attended TRFs, the ignored TRFs in the theta band showed a single main peak, M100_TRF_.

To quantify the differences between responses in the attended and ignored condition, individual time lags were extracted for each participant, listening scenario, and response type. The utilized search intervals for each frequency band are depicted in Table 2. For the broadband feature, a M100_TRF_ was found in 94 % of all participant-level TRFs, whereas a M200_TRF_ was present in 91 % of broadband TRFs. In the delta band, M100_TRF_ appeared in 84 %, M200_TRF_ in 91 %, and M400_TRF_ in 90 % of all participant-level TRFs. In the theta band, a M100_TRF_ was identified for every subject.

**Table 2:** Search intervals for the peaks in the subject-level TRF magnitudes for the different frequency bands.

| Frequency Band | M100<sub>TRF</sub> | M200<sub>TRF</sub> | M400<sub>TRF</sub> |
| --- | --- | --- | --- |
| Broadband | [80 ms, 140 ms] | [150 ms, 300 ms] | – |
| Delta Band | [20 ms, 160 ms] | [170 ms, 350 ms] | [360 ms, 710 ms] |
| Theta Band | [40 ms, 200 ms] | – | – |

The amplitude of M100_TRF_ was significantly higher in the attended than in the ignored condition in the broadband (*p* = 1e*−*9) and the delta band (*p* = 1e*−*12). In the theta band, however, no significant differences between attended and ignored response amplitude were found. The same applies to M200_TRF_ in the broadband TRFs, where no considerable differences in attended and ignored response amplitudes were discernible. In contrast, in the delta band, both M200_TRF_ and M400_TRF_ magnitudes were significantly greater in the attended TRFs compared to the ignored TRFs (M200_TRF_: *p* = 2e*−*5; M400_TRF_: *p* = 2e*−*8).

Furthermore, noticeable differences between attended and ignored responses were visible not only in the TRF magnitudes, but also in the time lags of the TRF peaks. Both M100_TRF_ and M200_TRF_ showed significant differences in delay, with the attended response occurring later than the ignored response in the broadband (M100_TRF_: *p* = 8e*−*3, M200_TRF_: *p* = 2e*−*11), the delta band (M100_TRF_: *p* = 0.01, M200_TRF_: *p* = 7e*−*5), and also the theta band (M100_TRF_: *p* = 5.0e*−*4). No significant differences in response peak time lags were found for M400_TRF_. These differences were also mostly reflected in the mean TRF peak time lags shown in Figure 3a, where the attended response peaks occurred with a delay of 10 ms to 40 ms compared to the ignored response peaks in all three frequency bands.

### Correlation Between Neural Responses and Participant Behavior

To investigate whether the neural responses were related to behavior, we computed Pearson’s correlation coefficient between the subject-level TRF magnitudes and the subject’s percentage of correctly answered comprehension questions (Figure 3b). For both the attended and the ignored listening mode, correlations were either close to zero, indicating that there was no relationship, or slightly positive ranging up to *r* = 0.25 (M100_TRF_ for attended audio input in the theta band). However, only the Pearson’s correlation coefficients estimated for the theta band were significant, revealing similar positive correlations for both ignored (*r* = 0.23*, p* = 0.04) and attended (*r* = 0.25*, p* = 0.02) modes.

There was no significant difference in the comprehension scores between musicians and non-musicians (*p* = 0.35).

### Cortical Responses in Musicians and Non-Musicians

To assess the putative influence of musical training on the cortical speech tracking, we compared the neural responses for the group of musicians to that of non-musicians (Figure 4). Similar to the population-level TRF magnitudes depicted in Figure 3a, for both groups attention-induced differences in the TRF magnitudes are visible in both the broadband and in the delta band. At the same time, no considerable differences between ignored and attended responses are discernible in the theta band.

**Figure 4:**
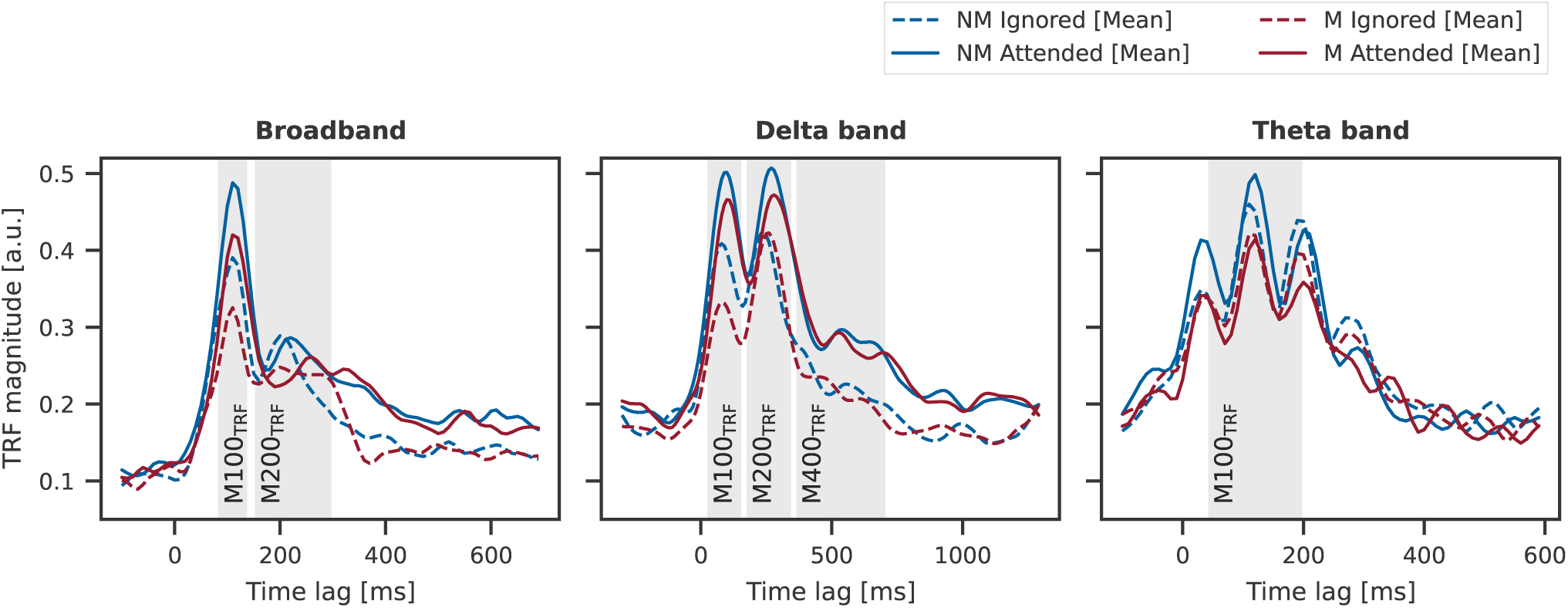
Magnitudes of the temporal response functions (TRFs) averaged across all sources for non-musicians (NM, blue) and musicians (M, red) in the attended condition (solid) and in the ignored case (dashed). The search areas for the subject-wise TRF peaks are shaded in grey. They were estimated based on the participants’ average signal and are thus identical for M and NM and ignored and attended responses. For all three frequency bands and both attended and ignored responses, a tendency towards higher M100_TRF_ peaks in non-musicians can be seen compared to the musicians.

For M100_TRF_ peaks across all frequency bands and listening modes, the mean magnitude was smaller in musicians than in non-musicians. This difference was most pronounced for the ignored M100_TRF_ response in the delta band with a mean increase of 23.0% in non-musicians compared to musicians. M200_TRF_ is also slightly larger for non-musicians than musicians in both the attended (9.6%) and ignored (16.5%) broadband TRF and the attended delta-band TRF (7.4%). For the remaining TRF peaks, no noticeable differences emerged. A difference in latencies could be observed in the attended M200_TRF_ in the broadband response, as the musicians’ average peaked 40 ms after the non-musicians at 260 ms. A similar effect was discernible in the delta-band M200_TRF_ in the ignored condition, where the musicians’ peak was delayed by 20 ms. Peak latencies were otherwise similar in both groups.

To assess the statistical significance of these differences, the subject-level TRF peaks and their time lags were compared between musicians and non-musicians. Furthermore, intra-group differences between ignored and attended response magnitudes characterized by the attentional modulation index *AI* were quantified. The results for all three frequency bands are depicted in Figure 5.

**Figure 5:**
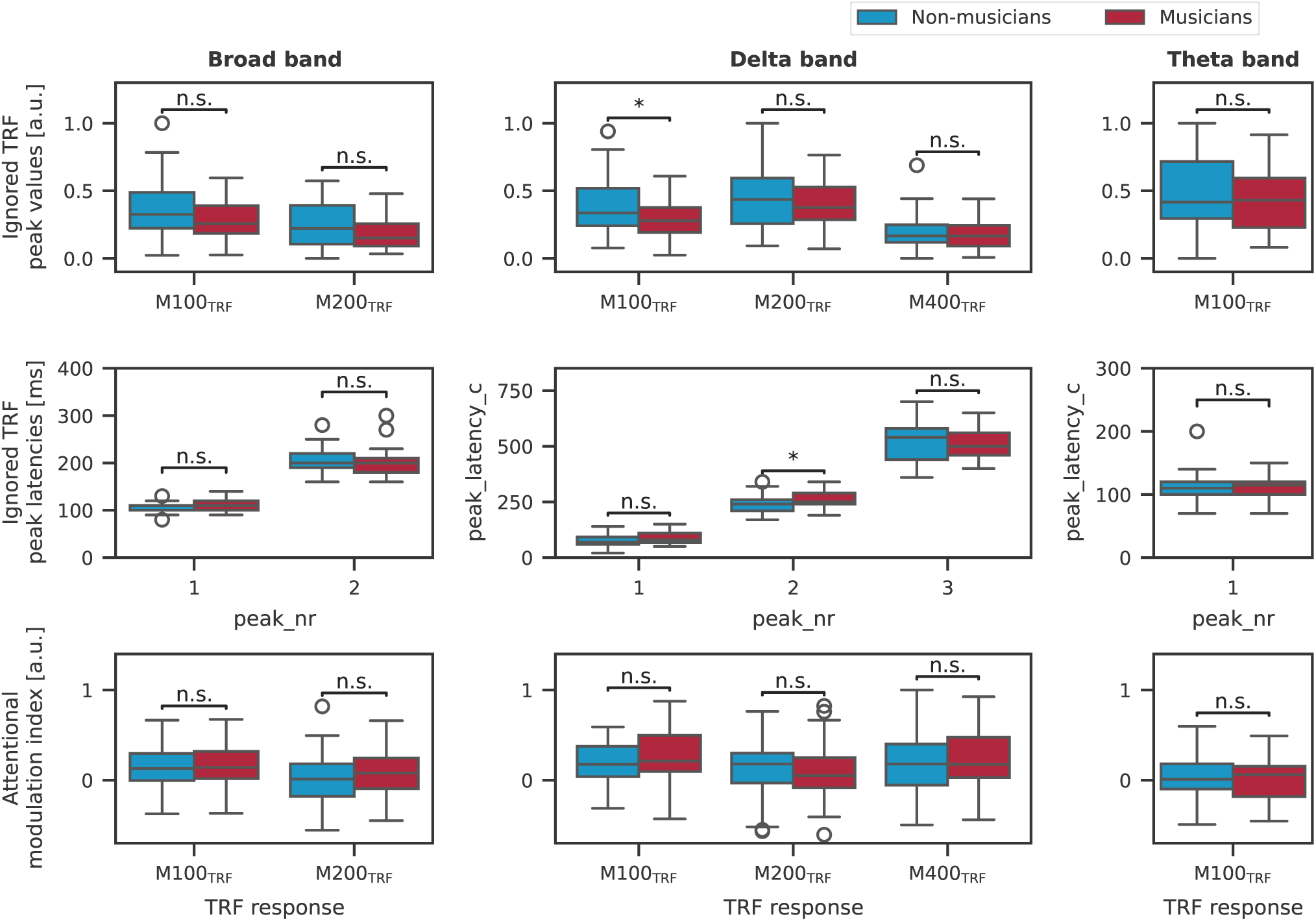
Comparison of subject-level TRF peak amplitudes (upper row), latencies (middle row) and attentional modulation indices (lower row) between musicians and non-musicians. For non-musicians, slight tendencies towards higher peak values in ignored M100_TRF_ and M200_TRF_ are recognizable, but only the ignored delta band M100_TRF_ is slightly significant (*p* = 0.049). For musicians, a significantly longer latency occurs for the M200_TRF_ in the ignored delta band. In turn, musicians show marginally larger attentional modulation indices for these response types. Peaks with significant differences in amplitude between the attended and ignored condition are marked with asterisks indicating their significance level (*, 0.01 *≤ p <* 0.05; **, 1e*−*3 *≤ p <* 0.01)

A tendency of increased M100_TRF_ amplitude in non-musicians compared to musicians was still observable. However, these differences were only slightly significant for the ignored delta M100_TRF_ (*p* = 0.049). All other disparities were not significant. The same applied to the latency of the M200_TRF_ responses, except for the latency of the M200_TRF_ in the delta band in the ignored condition, which was found to be significantly longer in musicians than non-musicians (*p* = 0.03). In turn, attentional modulation was highest at M100_TRF_, with musicians tending to show larger *AI* values than non-musicians in the broadband and the delta-band response. This difference was most pronounced in the delta band yielding an average attentional modulation index *AI* = 0.27 for musicians and *AI* = 0.20 for non-musicians. Additionally, in the broadband M200_TRF_, musicians showed a stronger representation of the attended feature yielding *AI* = 0.1 on average, whereas non-musicians had a modulation value of *AI* = *−*0.01, suggesting no modulation of attended and ignored cortical representations. However, none of these differences in the attentional modulation coefficient were statistically significant. Furthermore, in the theta band, the mean *AI* values of both groups were negligible (*AI* = 0.01).

### Relationship of Musicality Features and Cortical Responses

As a more fine-grained assessment of the influence of musical training on cortical speech tracking, we investigated whether there were relations between the subject’s different aspects of musical training and the TRF peak amplitudes.

The estimated contributions of each of the aspects of musical training to predicting the neural responses are depicted in Figure 6. No considerable contributions were found for the weekly practice feature. Feature weights for the number of instruments indicate a positive contribution of this property to the peak amplitudes throughout all frequency bands and listening scenarios, but did not reach statistical significance. Only two features, namely the starting age of learning an instrument and and the playing duration in years, were significant predictors. The former significantly contributed to the attended M200_TRF_ in the delta band (*p* = 0.01), with an assigned feature weight of 0.54, whereas the latter exhibited a negative relation with the TRF peak amplitudes. This observation became significant for the ignored M100_TRF_ in the delta band, where the smallest model coefficient was estimated (*−*0.55, *p* = 0.04).

**Figure 6:**
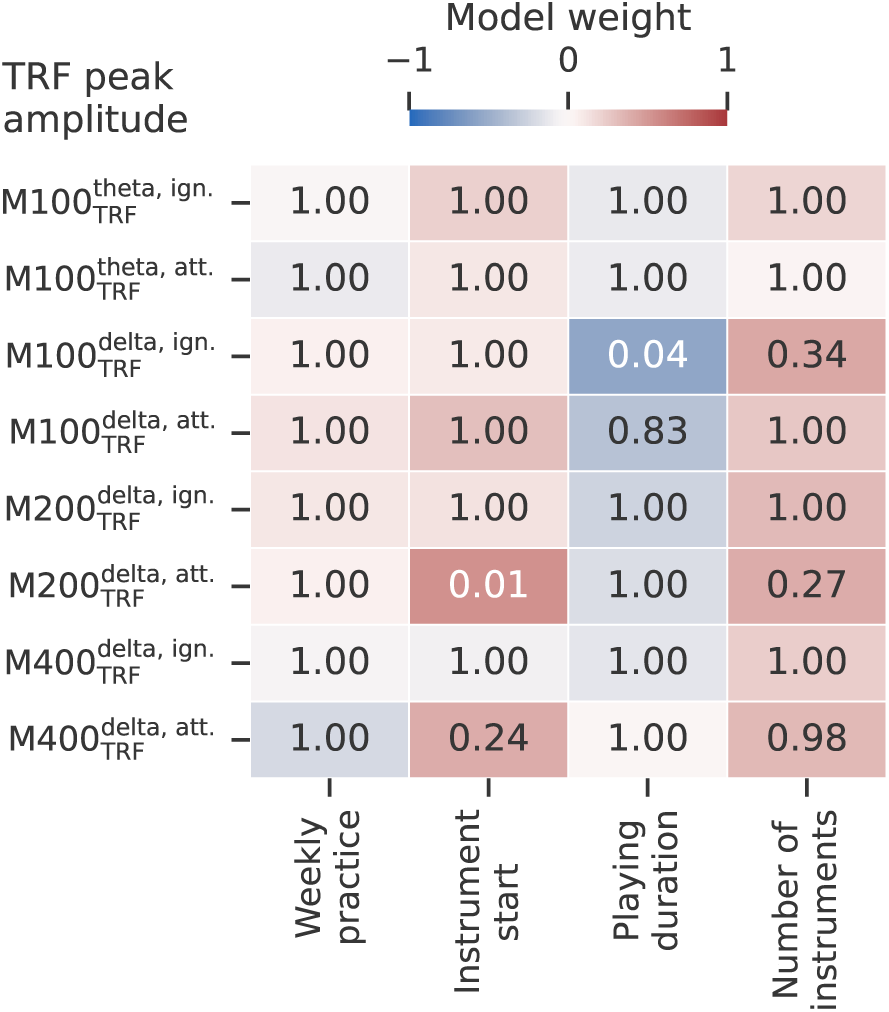
Influence of the individual aspects of musical training on the cortical responses. For each response type, frequency band, and attention mode, a multiple linear regression model was fitted to predict the response amplitude from the four aspects of musical training (hours of current weekly practice, starting age of playing an instrument, total duration of playing instruments, and the number of played instruments). The resulting weights of the linear model are assessed for statistical significance (*p*-value displayed in the corresponding cell). Two correlated aspects of musical training were found to be significant, the starting age and the duration of playing. Whereas the former is positively contributing to the attended M200_TRF_ response in the delta band (0.54), the latter is negatively affecting the ignored M100_TRF_ delta band response (*−*0.55).

## Discussion

In this work, we utilized a large dataset of MEG recordings from 52 participants, each of whom listened to 37 minutes of continuous speech. We thereby found that the attentional modulation of the cortical speech tracking occurred in the delta band but not in the theta band. We further found that musical training had only a slight influence on the neural responses, with most aspects of the latter remaining unaffected by musicianship.

### Different Types of Cortical Responses Masked in Broadband Become Apparent in the Delta and Theta Band

The speech envelope is a basic auditory feature that conveys various acoustic and linguistic information [14, 13]. The low-frequency range, below 20 Hz, is of particular interest, as it contains the rates of words and syllables.

Previous work has not only shown strong cortical envelope tracking in this frequency range [15] but also hypothesized that distinct functions can be attributed to delta- and theta-band tracking [25]. In particular, theta-band tracking is assumed to reflect lower-level syllable processing [50, 26, 23], whereas higher-level word processing is associated with delta-band responses [51, 52, 26].

Our results show noticeable differences between the cortical responses in the delta and theta bands, which remain obscured in the broadband due to the superposition of the individual components. Whereas in the theta band the most prominent response occurs around 100 ms, which is also visible in the delta band and broadband, additional response peaks are present in the delta band around 250 ms and 500 ms. Even though the latter is considerably weaker than the other response peaks, it is still clearly recognizable. In turn, in the broadband signal, the M200_TRF_ is discernible but has a much lower amplitude than in the delta band. No later response peak emerges in the broadband response.

### Low-Frequency Speech Envelope Tracking is Mainly Right-Lateralized

To obtain a better understanding of the underlying mechanisms of the observed cortical responses, we also looked at their spatial distribution and lateralization. Except for the late M400_TRF_, all cortical responses were significantly right-lateralized, with the highest activations centered in Heschl’s gyrus, i.e., the primary auditory cortex. This is in line with previous findings that also found a right-lateralization of low-frequency envelope tracking [53]. Furthermore, bottom-up processes on the sublexical level are dominated by the right hemisphere [54, 55]. For top-down semantic processing starting from the word level, however, a lateralization shift towards the left hemisphere has been observed in previous studies, which might cause the lack of lateralization of the M400_TRF_ observed here [24, 51].

### Delta and Theta Band Responses are Modulated Differently by Selective Attention

It is well-researched that in a competing speaker scenario, the envelopes of both speech streams are tracked individually in the auditory cortex, with the cortical response to the ignored speaker being weaker than to the attended speaker [1]. We recover this behavior in the broadband for the M100_TRF_ but not for the M200_TRF_. Because the broadband response contains both the delta and the theta band, the lack of attentional modulation of its M200_TRF_ peak is presumably due to the conflation of these two individual frequency bands.

Therefore, we focused on the separate assessment of the cortical tracking in the delta and theta bands. In the delta band, attended speech leads to significantly stronger tracking than ignored speech, across a large range of delays and encompassing the M100_TRF_, M200_TRF_, and M400_TRF_. In contrast, the theta band did not exhibit attentional modulation. However, we found a significant positive relationship between the theta band’s M100_TRF_ and the behavioral performance of the participants, which is in line with the crucial role of theta-band tracking in speech-in-noise intelligibility [22].

Because an M100_TRF_ emerges both in the delta- and in the theta-band tracking, but is only affected by attention for the delta-band response, this peak is likely not evoked by acoustic activity like an event-related potential. In fact, already the earliest evoked potentials in the cortex are modulated by attention [56], as opposed to the theta band’s M100_TRF_. Therefore, we conclude from our results that delta- and theta-band tracking of the envelope are continuously executed in parallel, whereby theta tracking provides bottom-up information about the unfiltered auditory input, while in the delta band, background information is attenuated and further processed for linguistic information.

A time delay of around 10 ms between the peaks in the responses to ignored and attended speech was reported in previous work [1], with the attended response peak occurring earlier than the ignored response peak. However, in our study, we observed the opposite effect: the response peaks of the ignored stream appeared significantly before the response peaks of the attended stream. This could indicate that the ignored stream is processed less thoroughly than the attended one, and therefore the response arises sooner.

### Few Significant Differences between Musicians and Non-Musicians

When assessing the influence of musical training on cortical speech tracking, we only found minor significant effects on the M100_TRF_ and M200_TRF_ in the delta band. For the ignored speech signal, the delta band M100_TRF_ peak was slightly greater in non-musicians than in musicians. Furthermore, its magnitude was negatively impacted by the duration over which participants had played an instrument. Furthermore, the ignored M200_TRF_ peak was slightly delayed in musicians than in non-musicians. Additionally, its magnitude resulting from the attended speech signal had a positive correlation with the starting age of practicing an instrument. The other components of the neural speech tracking were not significantly affected by musicianship, although we observed a general trend of lower amplitudes for musicians as compared to non-musicians.

To interpret these findings, we note that, in contrast to intra-subject modulation effects, where stronger tracking of a speech signal in background noise is indicative of better intelligibility, this does not necessarily apply when comparing cortical responses *between* participants. Previous studies investigating speech perception in the elderly in fact demonstrated that stronger cortical speech tracking correlated with lower intelligibility scores [57, 58]. The strength of the neural tracking might thus not indicate improved comprehension, but rather increased listening effort.

In line with these deliberations, Jantzen et al. (2014) observed that non-musicians showed increased cortical activity, especially for early responses [33]. This finding matches with the tendencies towards higher response magnitudes in non-musicians, which we observed here (despite a lack of statistical significance). At the same time, this might explain the slightly higher attentional modulation indices for early delta responses in musicians.

This ‘inverse’ relationship between musicianship and the strength of the cortical tracking is also reflected in the relation between the aspects of musical training and the neural responses that we determined. The earlier the participants started playing an instrument, the weaker the delta-band responses were. This aspect of musical training hence seems to have a strong influence on speech-in-noise perception, which is consistent with the results of other studies [2, 8].

We expect the M400_TRF_ to be most closely associated with semantic integration and contextualization of the speech signals. This response was remarkably similar between musicians and non-musicians. Together with the insignificance of most of the other aspects of the neural speech tracking that we examined, we conclude that cortical speech tracking is essentially not affected by musical training. The behavioral improvements in speech-in-noise perception observed in musicians appear to originate in other neural mechanisms, probably further downstream of the cortical speech tracking since the latter already involves early auditory responses.

## Conclusions

In summary, our study showed that the attentional modulation of cortical speech tracking results from the delta band but not from the theta band. The theta band presumably reflects lower-level acoustic processing such as syllable parsing, which appears to be equally executed for the attended and the ignored speech stream. Neural activity in the delta band, in contrast, is responsible for higher-level linguistic processing, and our results show that attentional effects emerge at this stage. Musical training leaves both types of responses largely unchanged, even though we observed a tendency towards stronger cortical speech tracking in people with less musical training.

## Data availability

The MEG data used in this study is available from the authors upon request.

## Author contributions

**Alina Schüller, Annika Mücke**: Conceptualization, Interpretation of Results of Experiments, Formal analysis, Investigation, Writing original draft. **Jasmin Riegel**: Conceptualization, Data Acquisition, Reviewing and Editing of the paper. **Tobias Reichenbach**: Conceptualization, Interpretation of Results of Experiments, Supervision, Reviewing and Editing of the paper.

## Acknowledgments

This project was supported by the German Federal Ministry of Education and Research (Cluster4Future SEMECO, project number 03ZU1210FB) and the German Science Foundation (DFG, project number 523344822).

## Conflict of interest statement

The authors declare no competing financial interests.

## Notes

### Competing Interest Statement

The authors have declared no competing interest.

